# Automated segmentation of synchrotron-scanned fossils

**DOI:** 10.1101/2024.10.23.619778

**Authors:** Melanie A. D. During, Jordan K. Matelsky, Fredrik K. Gustafsson, Dennis F. A. E. Voeten, Donglei Chen, Brock A. Wester, Konrad P. Körding, Per E. Ahlberg, Thomas B. Schön

## Abstract

Computed tomography has revolutionised the study of the internal three-dimensional structure of fossils. Historically, fossils typically spent years in preparation to be freed from the enclosing rock. Now, X-ray and synchrotron tomography reveal structure that is otherwise invisible and data acquisition can be fast. However, manual segmentation of these 3D volumes can still take months to years. This is especially challenging for resource-poor teams, as scanning may be free, but the computing power and (AI-assisted) segmentation software required to handle the resulting large data sets are complex to use and expensive.

Here we present a free, browser-based segmentation tool that reduces computational overhead by splitting volumes into small chunks, allowing processing on low-memory, inexpensive hardware. Our tool also speeds up collaborative ground-truth generation and 3D visualization, all in-browser. We developed and evaluated our pipeline on various open-data scans of differing contrast, resolution, textural complexity, and size. Our tool successfully isolated the *Thrinaxodon* and *Broomistega* pair from an Early Triassic burrow. It isolated cranial bones from the Cretaceous acipenseriform *Parapsephurus willybemisi* on both 45.53 *µ*m and 13.67 *µ*m resolution scanning data. We also isolated bones of the Middle Triassic sauropterygian *Nothosaurus* and a challenging scan of a squamate embryo inside an egg dating back to the Early Cretaceous. Our tool reliably reproduces expert-supervised segmentation at a fraction of the time and cost, offering greater accessibility than existing tools. Beyond the online tool, all our code is open source, enabling contributions from the palaeontology community to further this emerging machine learning ecosystem.

## I. Introduction

Fossilisation is rare. Biological remains require very specific circumstances to be preserved over a long period of time. Subsequently, these fossils need to be found and appropriately extracted for academic study. Physical preparation of fossils used to be the only way to explore fossil contents. Yet preparation is not without risks, as it can damage the bone surface, and it will remove potential soft tissues that are not always recognisable upon exposure. Furthermore, it is not guaranteed to reveal all of the anatomy of interest. Museum curators are understandably hesitant to allow for destructive analyses or extensive preparation of delicate structures. Recent technological advances, such as Computed Tomography (CT) in general and Propagation Phase-Contrast Synchrotron Radiation Micro-Computed Tomography (PPC-SR*µ*CT) [1] in particular, have made it possible to study fossils at sub-micron resolutions, revealing parts of the anatomy that were previously hidden by matrix, and enabling the study of internal anatomical features or preserved soft tissue without damaging the fossils. This innovation has dramatically increased our ability to interrogate otherwise impenetrable palaeontological mysteries.

Despite these advantages, the 3D volumes produced by X-ray tomography (XRT) require substantial digital storage and computing power. The subsequent segmentation and modelling workflows thus tend to demand substantial funds not universally available to research groups and can take so much manual labour that projects remain unfinished. Commercial endeavours offer machine learning-based segmentation as a solution to these challenges and have already made a significant impact in the field [2], but these tools are typically designed for industrial applications and are therefore fine-tuned for neither synchrotron nor palaeontological data. To meet the need for palaeontology-focused, accessible, and democratised XRT analysis software, our team developed an online segmentation pipeline for fossils, trained on publicly available synchrotron data, and offered as a free service to the public **(Fig. 1)**. Here, we present the *ml4paleo* software suite, a Python package that combines traditional machine learning tools, neural image segmentation tools, and batch processing tools to segment large-scale palaeontological XRT data volumes. We share a simple “online-learning” web interface for machine-guided, 2D slice-based image annotation to aid in training small models without downloading new software. Our solution can be operated with minimal technical expertise and runs on commodity computing hardware. Our codebase is open source and we encourage community feedback and contributions.

**Fig. 1.**
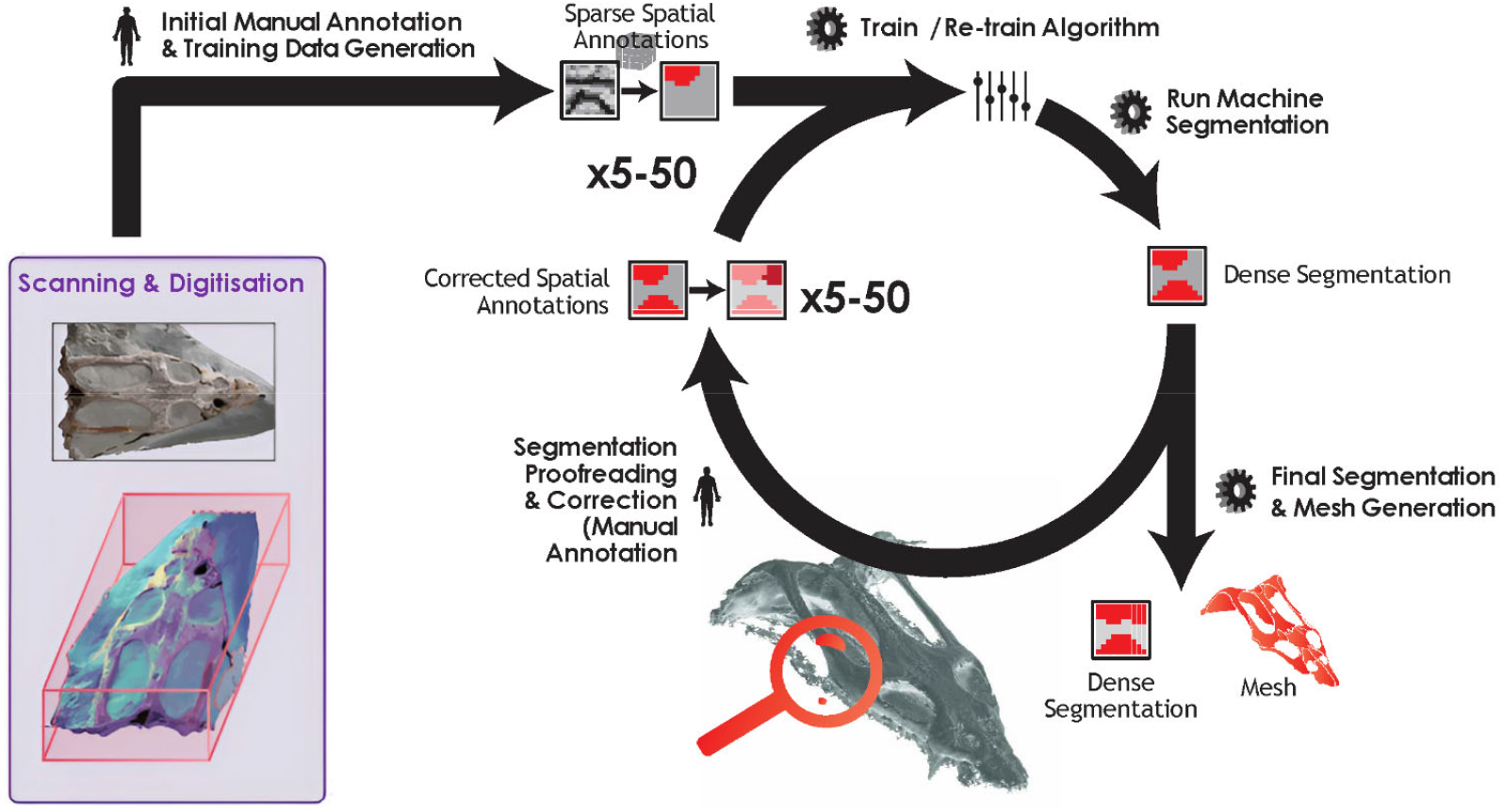
An overview of the pipeline proposed here. Raw X-ray image volumes (left) are iteratively segmented through collaborative human-machine teaming. Machine-guided annotation in our web application yields training data for the segmentation models. The segmentation model is applied to the whole dataset through batch processing, taking advantage of chunked data storage paradigms. This process can be repeated until the model meets human-defined quality benchmarks. Finally, a high-resolution segmentation mask can be exported, alongside image renders and meshes suitable for rendering in 3D software.

**Fig. 2.**
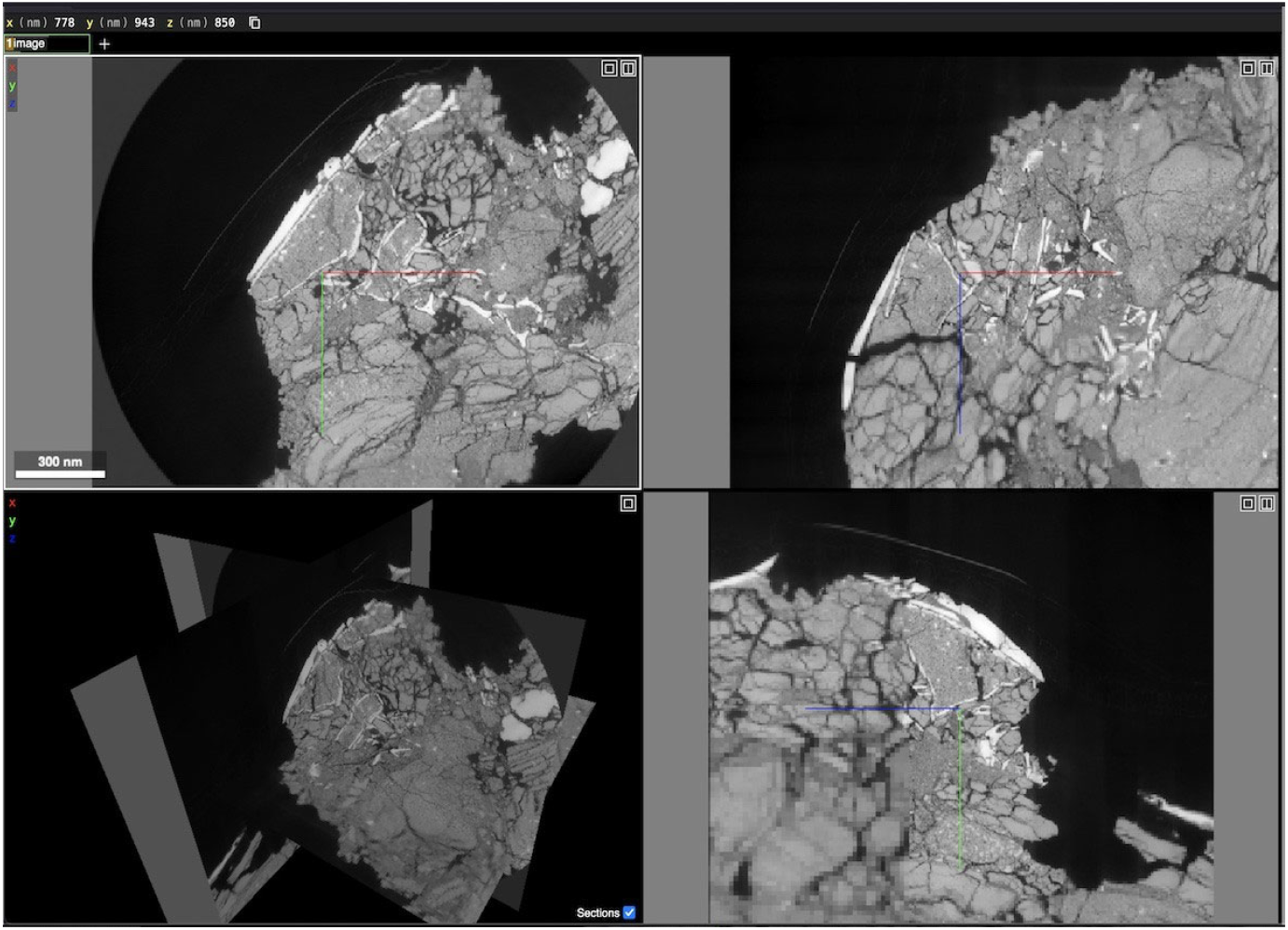
Browser-based visualisation. This stage in the workflow uses *Neuroglancer* to render imagery and annotations. This figure shows a user visualising the *Parapsephurus willybemisi* 13.67 *µ*m resolution scan [5] in interactive 3D in a modern web browser. This tool is included with the *ml4paleo* web suite.

**Fig. 3.**
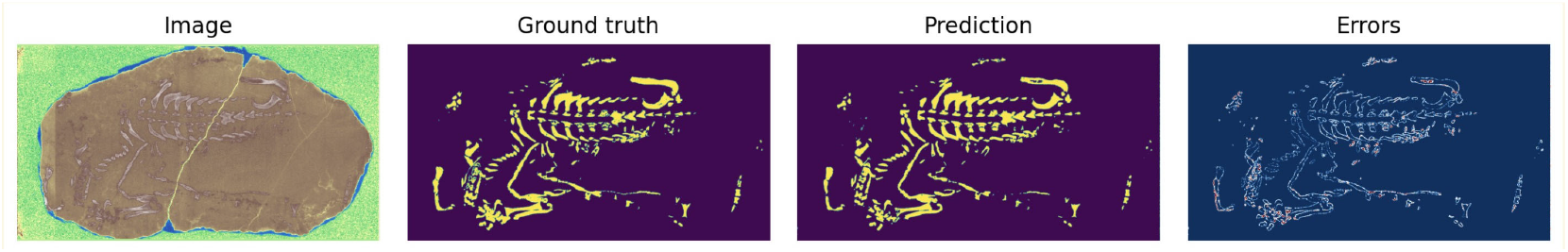
Performance on the burrow dataset. We picture one 2D slice of the original volume (left, artificial color), as well as human and machine annotated segmentation masks. On the far right, we render the residuals.

Automated segmentation is a major ongoing challenge across the XRT community broadly [3, 4]. The specific challenges in palaeontology, such as the variation in contrast through an image stack, and high absorption caused by metallic inclusions, reflect similar challenges that other fields face with their own XRT workloads. We therefore intend for our tool to serve as a model and inspiration for a wide range of fields, such as neuroscience, histology, and earth sciences.

## II. Materials and Methods

### A. Materials

We leveraged five public datasets that are available online in the *paleo*.*esrf*.*eu* database, a public resource for palaeontological volumes scanned at the European Synchrotron Radiation Facility (ESRF) [6].

The datasets used in this study represent a range of fossil types and scanning conditions, each presenting its unique challenges to segmentation. The Burrow dataset, from the Early Triassic Karoo Basin in South Africa, bears a *Thrinaxodon* and a *Broomistega* and was scanned at 45.5 *µ*m resolution [7]. This dataset benefits from a relatively clear contrast between the fossils and the surrounding matrix, allowing for high segmentation accuracy. In contrast, the Middle Triassic *Nothosaurus* skull from the Muschelkalk of the Netherlands scanned at 12.82 *µ*m resolution [8] presents lower contrast. This makes it more difficult to distinguish between fossilised bone and matrix. The *Nothosaurus* stack was also post-processed into a recoded version of the three-dimensional data based on local texture complexity to reveal low-contrast features [8]. The end-Cretaceous paddlefish *Parapsephurus willybemisi*, from the Tanis deposit in North Dakota, was scanned at 45.53 *µ*m and 13.67*µ*m resolution [5, 9]. These datasets highlight the role of scan resolution in segmentation accuracy. The higher-resolution ∼13µm dataset allows for the segmentation of more detailed phenomena, while the lower-resolution ∼45µm dataset does not, as key anatomical features become more difficult to discern.

Finally, an Early-Cretaceous squamate embryo inside an egg from Thailand (Phu Phok) was scanned at 5.06 *µ*m resolution [10]. This dataset presents a unique challenge due to the fine structures, compounded by low contrast, leading to a significant challenge in segmentation for both humans and machine learning-based methods. For additional details about the subsamples of these datasets used for our evaluation, see **Table I**.

**TABLE I.** Datasets used in this study.

| Dataset nickname | Taxa | Age | Location | Dataset Size | Resolution | Reference |
| --- | --- | --- | --- | --- | --- | --- |
| Paddlefish ~45um | *Parapsephurus willybemisi* | Cretaceous | Tanis Deposit (ND, USA) | 1771, 1117, 200 | 45.53 $\mu\text{m}$ | [5] |
| Paddlefish ~13um | *Parapsephurus willybemisi* | Cretaceous | Tanis Deposit (ND, USA) | 3815, 3815, 200 | 13.67 $\mu\text{m}$ | [5] |
| Burrow | *Thrinaxodon, Broomistega* | Triassic | Karoo Basin (South Africa) | 1685, 3043, 200 | 45.5 $\mu\text{m}$ | [7] |
| Phu Phok | Anguimorph embryo | Early-Cretaceous | Phu Phok (Thailand) | 1939, 2206, 200 | 5.06 $\mu\text{m}$ | [10] |
| Nothosaur | *Nothosaurus marchicus* | Middle-Triassic | Winterswijk (The Netherlands) | 1410, 1410, 200 | 12.82 $\mu\text{m}$ | [8] |

### B. Web application architecture

The *ml4paleo* web application comprises four components; one main application server, and three task runners that operate on an on-disk queue.

#### 1) Application server

The web application server is written in Flask [11]. The server can use an on-disk serialisation format for storing dataset metadata, or it can be configured to run a standalone database daemon. By default, our application uses a standalone JSON file as a metadata store to reduce the software dependencies of running a minimal deploy.

#### 2) Conversion queue

Researchers commonly receive concatenated image stacks as output from the synchrotron facility. This image stack tends to be too large and inefficient to access for common machine learning workflows. Thus, the first memory-intensive stage of processing is to convert this inefficient data format into a chunked data format. The user can easily upload their dataset in most common formats, including image stacks and some common chunked formats. Chunking high-dimensional data involves re-organising a large dataset of 2D images into smaller, more compact 3D (volumetric) or 4D (volumetric time series) pieces, known as chunks. These chunks can then be treated as independent small volumes, or manipulated in spatial sequence. This conversion enables the *ml4paleo* tools to operate on sub-volume cutouts without loading complete slices into memory. It is this memory efficiency that enables our infrastructure to work equally well on low-performance and high-grade computers alike.

### C. Model architectures

As a core design principle for this software package, we aimed to select default algorithms that impose minimal restrictions on user hardware, ensuring accessibility for a wide range of systems. In other words, our goal was to accommodate the lowest common denominator in terms of hardware capabilities. This required that our segmentation algorithms be compatible with consumer-grade hardware while operating at the necessary spatial scales.

Two primary decisions emerged from this requirement: first, the dimensionality of the segmentation process, and second, the selection of machine learning models, based on both computational demands and dimensionality considerations.

While 2D stacked segmentation, where individual slices are segmented and later reconstituted into 3D, can deliver acceptable results in many cases, it is inferior to true 3D segmentation. The latter benefits from the inclusion of additional spatial context within the 3D data stack. However, memory limitations significantly reduce the efficiency of 3D segmentation, as its computational complexity scales with *N* ^3^ compared to *N* ^2^ for 2D segmentation. Given that X-ray tomography (XRT) data is isotropic, with uniform voxel sizes across all dimensions, 2D models have proven sufficient for our purposes thus far. Nevertheless, our codebase includes 3D models, allowing users to switch between 2D and 3D approaches as needed (see Data & Code Availability).

As the default segmentation algorithm, we utilise a simple Random Forest (RF) model trained on heuristic-based pixel features [12]. This is implemented using the *skimage*.*feature*.*multiscale basic features* module with default parameters, all of which are adjustable through the administrator settings of the web application. The feature extractor relies on basic image properties such as texture, intensity, and edge detection, progressively applying Gaussian filters to estimate these features on “downsampled” versions of the images.

For training efficiency, we optionally provide a parameter to subsample pixels, using every *n*th pixel for training rather than all annotated pixels. By default, 50% of the available image/segmentation pairs are used during the training process.

Our codebase also includes a convolutional residual U-Net [13], which requires a GPU for efficient training, though it otherwise operates with modest computational requirements. These tools are implemented through the *ml4paleo*.*segmentation*.*Segmenter3D* interface. Both this and the *ml4paleo*.*segmentation*.*Segmenter2D* interface are designed to support easy integration of more advanced or specialised segmentation algorithms by third-party contributors

### D. Data processing pipeline

In order to process arbitrarily large volumes of 3D imagery, it is necessary to decrease the size of the volume that is fed into the pipeline. In volumetric data processing, this is referred to as *chunking* the volume into subvolumes with contiguous areas nearby in memory. Several data standards have been built in a variety of scientific domains to meet this need [3, 14, 15]. For most purposes, interchangeable data standards make the distinctions between these chunked formats insignificant, and for this project we have used the standard *zarr* format [15]. This chunking technique makes it possible to independently segment subvolumes of the dataset, which can then be combined in a segmentation fusion step later. In their corresponding *Python* Application Programming Interfaces (APIs), these “datastores” tend to meet the standard numpy-like slicing interface in order to be maximally interoperable with the machine learning ecosystem [16]. This also means that data can be lazily loaded from disk, so that the majority of the volume can remain compressed and at rest while subvolumes are accessed in memory.

The fusion step can be quite nuanced: in some cases, a single pixel may be justifiably assigned to more than one segmentation type, and therefore it is generally advised to store segmentation as a multi-channel output. While this enables higher-quality segmentation, it comes at the cost of greatly increased storage space, as each segmentation mask must be stored independently. Because our intention was to minimise the computational and storage costs for new projects, we opt here to store single-channel segmentation by default, though this is configurable by the administrator.

### E. Training of segmentation model

We provide a simple web application to generate human expert ground-truth to train the segmentation pipeline. This tool (**Fig. 4**) provides a resizable brush for short annotation tasks of small 2D slices. The user can adjust the contrast and scroll several slices up and down in the stack to provide spatial context during annotation. The tasks are designed to be very short in duration to avoid attentional errors and to make exhaustive annotation feasible on resource-constrained hardware, following design principles from [17].

**Fig. 4.**
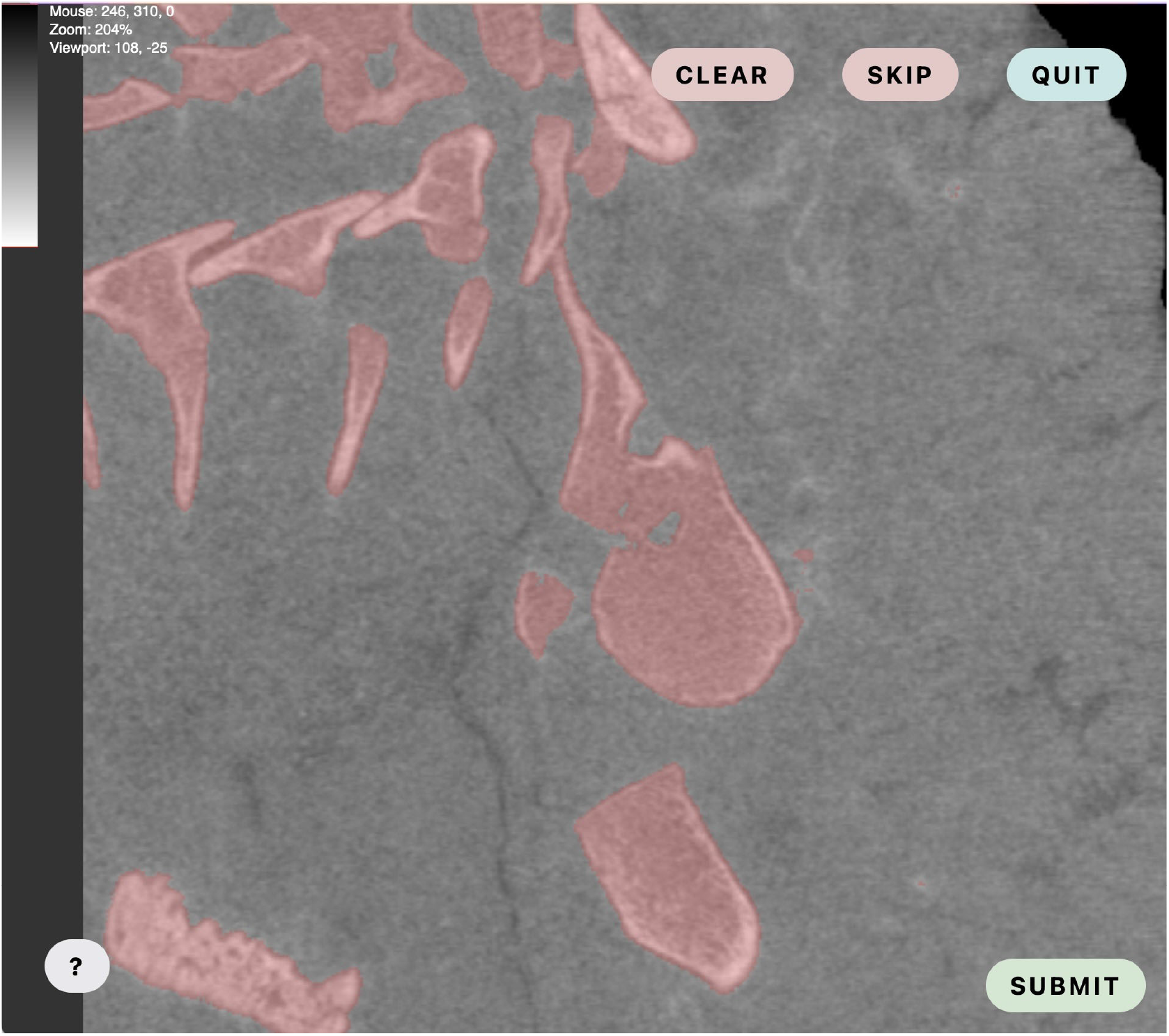
A screenshot of the annotation tool on an excerpt slice of the Burrow dataset. The user is given a resizable brush to manually segment fossils from the matrix in a small 2D area. Here, shown in red, the *ml4paleo* automated segmentation has already been run on this slice, so the user only needs to proofread the machine’s work, by adding with a brush or removing with an eraser, rather than annotate *de novo*.

Pairs of human-annotated segmentation masks and the original corresponding imagery are then fed to the selected machine learning model for training. By default, *ml4paleo* uses a random forest with local pixel-neighbourhood features [12], though the segmentation algorithm in use can be replaced by a simple U-net neural network [13] or other machine learning models that implement our *ml4paleo*.*segmentation*.*Segmenter3D* interface. A checkpoint of the trained model is then saved for reuse under a unique identifier that contains both the timestamp of training and the specific training samples used. This means that as more data is generated, a library of models can be accumulated to enable analysis and understanding of the relationship between training data volume and segmentation quality.

#### 1) Segmentation queue

Once a machine learning model has been trained for image segmentation, the dataset is queued for segmentation. In this stage, each worker loads a subvolume of the imagery, performs a dense segmentation, and saves a new segmentation mask volume to disk.

This new layer is identified by the unique model identifier that was generated during the model training; thus, segmentation layers can be uniquely associated with the model that generated them, for provenance and reproducibility.

#### 2) Mesh queue

As a postprocessing stage after segmentation, a user may opt to download their dataset as a mesh in OBJ or STL formats for rendering in 3D software. This mesh process also operates on one chunk at a time, after which the sub-meshes are stitched into one large mesh.

#### 3) Visualisation in three dimensions

To visualise the scan as well as the annotations and segmentations, we utilise *Neuroglancer*, a high-performance, WebGL-based visualisation framework developed by the Google Connectomics Team, designed for efficiently viewing and analysing large-scale volumetric neuroscience datasets [18]. A deployment of Neuroglancer can be packaged with deployments of the *ml4paleo* software suite, or any public Neuroglancer application can be used and pointed at ml4paleo data volumes through our volumetric API.

### F. Online learning

Once the user has produced at least one saved model — requiring a minimum of one annotated slice — they can optionally allow the machine to produce an initial hypothesis segmentation before each subsequent annotation task. This has the potential to greatly reduce the labour cost of individual annotation tasks, since the machine may produce a nearly-complete segmentation mask, depending on the image properties of the underlying volume. Furthermore, the available eraser tool enables users to refine segmented slices, significantly reducing the effort compared to re-segmenting from scratch. Through this machine guidance, the user will become aware of the qualitative performance characteristics of the segmentation model and can monitor its improvement throughout the annotation process.

In our approach, for the datasets tested in this study, a total of 20 slices (from patches of full slides) were initially annotated by the same expert paleontology segmentation author (MD) for each dataset using the annotation web application. However, due to the specific challenges posed by the Phu Phok and Nothosaur datasets, additional annotations were required to improve to the required level of performance. These properties and improvements are reported below.

Fossil embryos, such as those in the Phu Phok dataset, are typically poorly ossified [10], resulting in significant contrast variations between bones, soft tissues, and matrix. This contrast variability makes it very challenging for the human eye to differentiate structures. The same difficulty is encountered by automated segmentation systems, which struggle with this inconsistent presentation of biologically similar structures. Likewise, the Nothosaur dataset contained a crack running through the fossil’s skull, which created areas of phase contrast that obscured the underlying bone texture. Cracks like these are common in fossil scans and pose difficulties for both human and AI segmentation efforts.

Due to these challenges, after the initial 20 slices were annotated for both datasets, an additional 10 slices were segmented and the model was retrained. A final set of 5 slices was then annotated, resulting in a total of 35 annotated slices for the Nothosaur and Phu Phok Egg datasets to achieve satisfactory segmentation results. In our experiments, switching the default segmentation tool from the random forest to a more sophisticated (neural) model resolved many of these failures, at the cost of additional execution time and GPU requirements.

After the user is satisfied with the model performance, they can deploy the segmentation model on the complete data volume. Below, we provide quantitative evaluation of the *ml4paleo* web application default segmentation, showing performance metrics for leave-one-out cross-validated pairs of human-annotated segmentation and imagery.

### G. Chunked processing

Because *ml4paleo* datasets are stored in a chunked volumetric data format, it is easy to parallelise an operation across the dataset in small sub-volume increments. Using dataset slicing code borrowed from the electron microscopy neuroscience community [14], we run the latest model checkpoint from the training stage on each sub-volume of data. When processed in memory, the sub-volumes do not have to match the default size of the chunked format sub-volumes on disk (128 x 128 x 128). Our informal testing suggests that using a larger segmentation sub-volume size of 256 x 256 x 256 can significantly improve throughput without compromising segmentation quality. However, this depends heavily on the available computing resources: memory-constrained environments may benefit from smaller segmentation chunk sizes, while environments with limited CPU cores may perform better with larger chunk sizes.

It is beneficial to have a small overlap of the processed chunks so that the computed pixel-wise features on the edges of each chunk accurately reflect their spatial context (i.e. in the “middle” of the volume, not on an edge). However, we discovered in our tests that this was not always necessary, especially in more easy-to-segment datasets. Samples with a clear brightness contrast between matrix and fossil tended not to need additional context, and fossils distinguished from the matrix only by texture tended to need a broader margin of additional context from neighbouring chunks. A further evaluation and benchmark of this dataset property is left for future work.

### H. Evaluation Metrics

To better understand the segmentation performance, we compared it qualitatively to prior segmentations by the original researchers [5, 7, 8, 10] and quantitatively with our own cross-validation evaluation suite.

We report the following:

- **Precision** measures the proportion of correctly identified fossil pixels relative to all pixels predicted as fossil by the model. High precision indicates that the model made fewer false positive predictions.
- **Recall** assesses the model’s ability to identify all actual fossil pixels within the dataset. A higher recall means that the model successfully identified more true fossil pixels.
- **F1-score (Dice coefficient)** is the harmonic mean of precision and recall, providing a balanced measure of segmentation quality. It is particularly useful when precision and recall values are unbalanced.

## III. ResultS

The model’s performance on all five datasets can be found in **Table II** and reported visually in **Fig. 5**.

**TABLE II.** Performance of our default random forest pixel classifier on a variety of datasets. Values WERE COMPUTED USING LEAVE-ONE-OUT CROSS-VALIDATION WITH 2D SLICE TRAINING.

| Dataset | Precision | Recall | F1 |
| --- | --- | --- | --- |
| Burrow | $0.861 \pm 0.142$ | $0.726 \pm 0.154$ | $0.777 \pm 0.129$ |
| Nothosaurus | $0.602 \pm 0.227$ | $0.496 \pm 0.264$ | $0.522 \pm 0.24$ |
| Paddlefish ~13um | $0.583 \pm 0.298$ | $0.492 \pm 0.278$ | $0.488 \pm 0.257$ |
| Paddlefish ~45um | $0.718 \pm 0.427$ | $0.191 \pm 0.187$ | $0.185 \pm 0.181$ |
| Phu phok | $0.401 \pm 0.295$ | $0.02 \pm 0.014$ | $0.035 \pm 0.027$ |

**Fig. 5.**
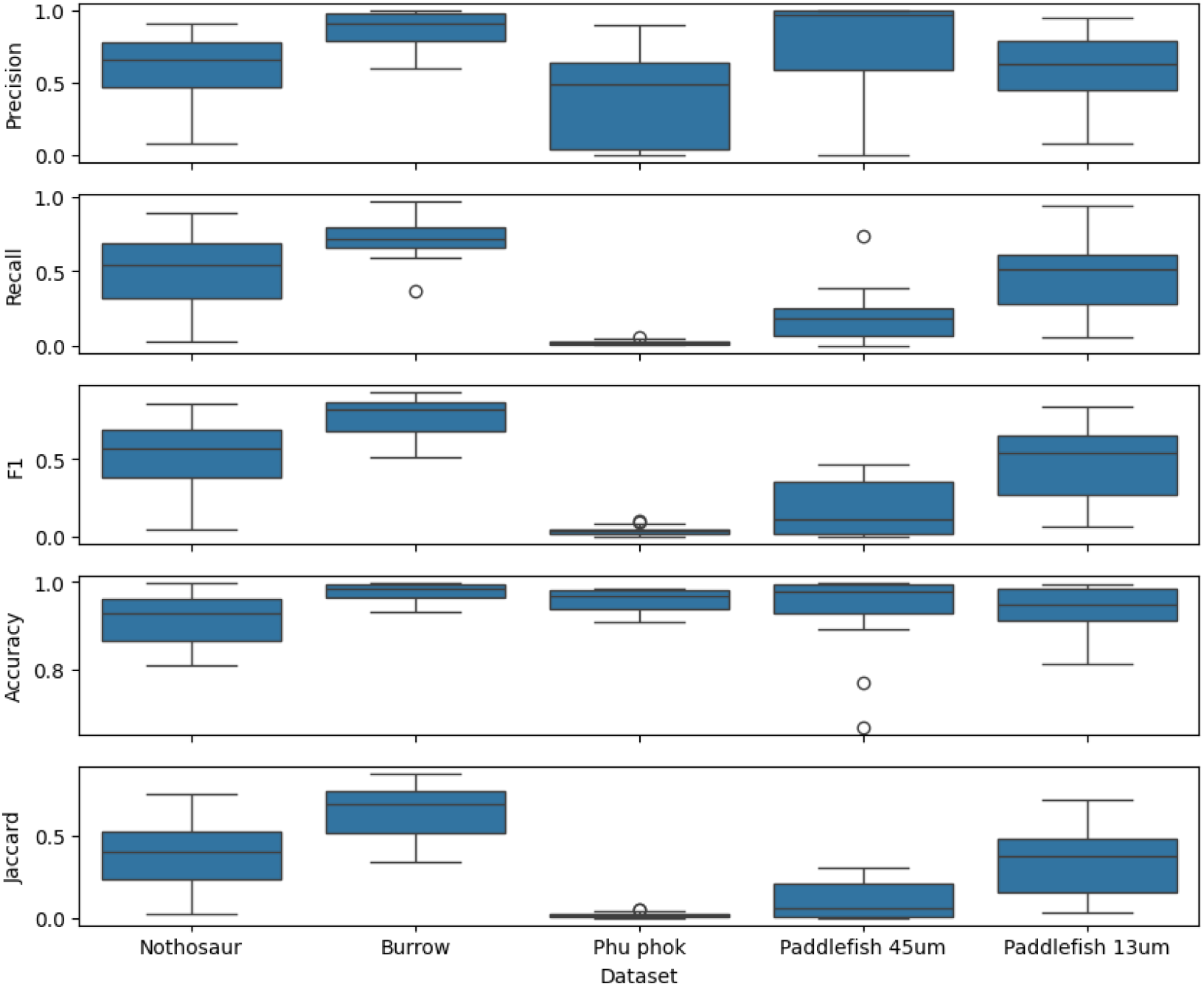
Performance metrics. An illustration of the major differences in segmentation quality between the different datasets, generally due to the inherent difficulty of distinguishing fossil pixels visually (i.e., through contrast or texture). Values are displayed in Table II.

### A. Burrow data, compared to manual segmentation

The segmentation model demonstrated strong performance, with high precision and recall, suggesting a robust ability to correctly identify and segment relevant features. The accuracy was also high (0.982), while the Jaccard Index, a measure of similarity between predicted and true masks, was 0.759, indicating a solid overlap between machine predictions and human annotations (**Fig. 6**).

**Fig. 6.**
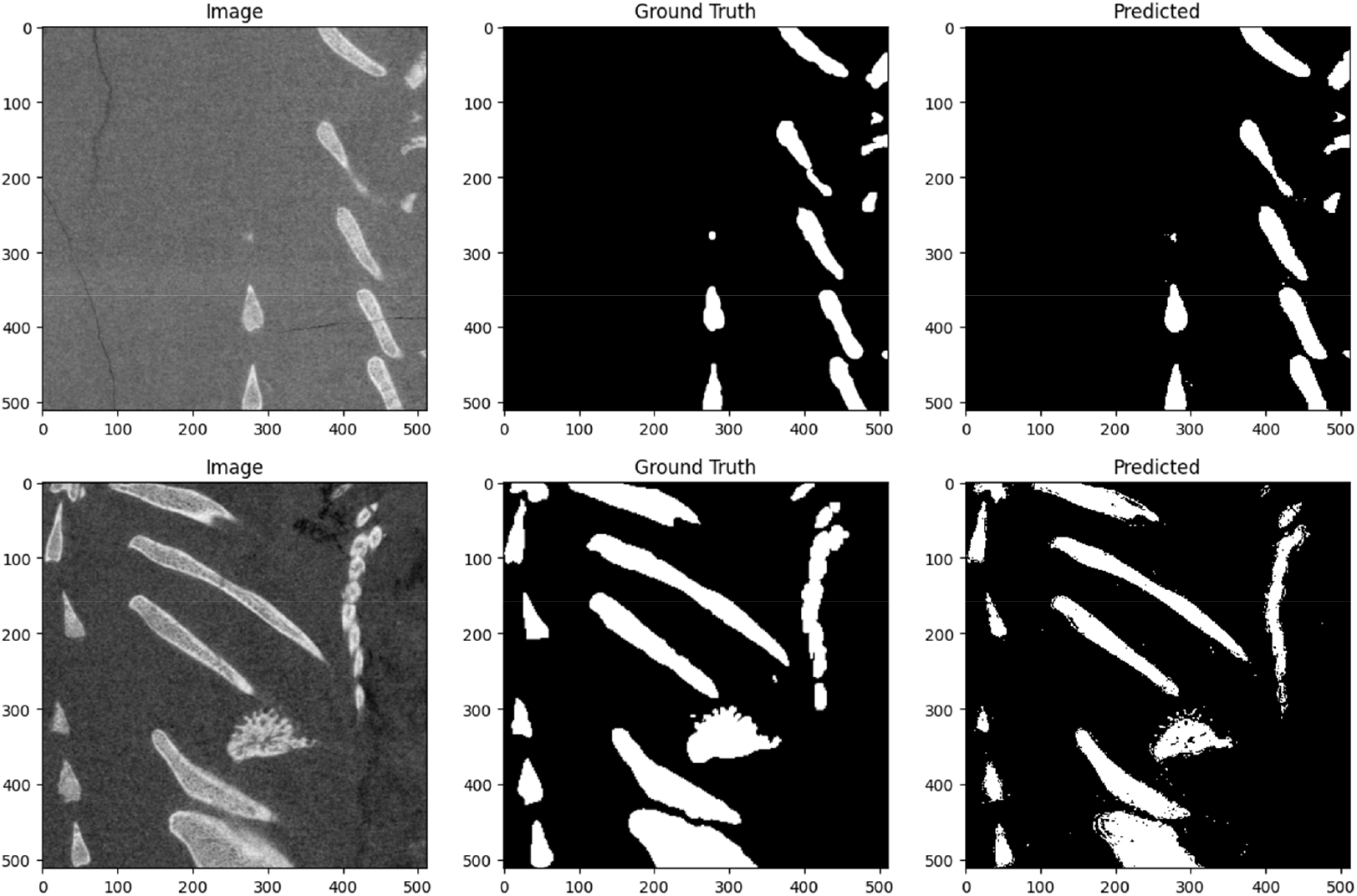
Burrow segmentation samples. The high contrast of the fossil versus the matrix was well suited for the simple multiscale pixel features used by our default model.

### B. Paddlefish 13.67µm and 45.53µm scans, compared to manual segmentation

Paddlefish ∼13*µ*m Dataset (**Fig. 7**): For this high-resolution scan, performance was moderate, with precision (0.583) and recall (0.492), resulting in an F1-score of 0.488. The segmentation shows a reasonable but not perfect overlap between predicted and true segmentation masks.

**Fig. 7.**
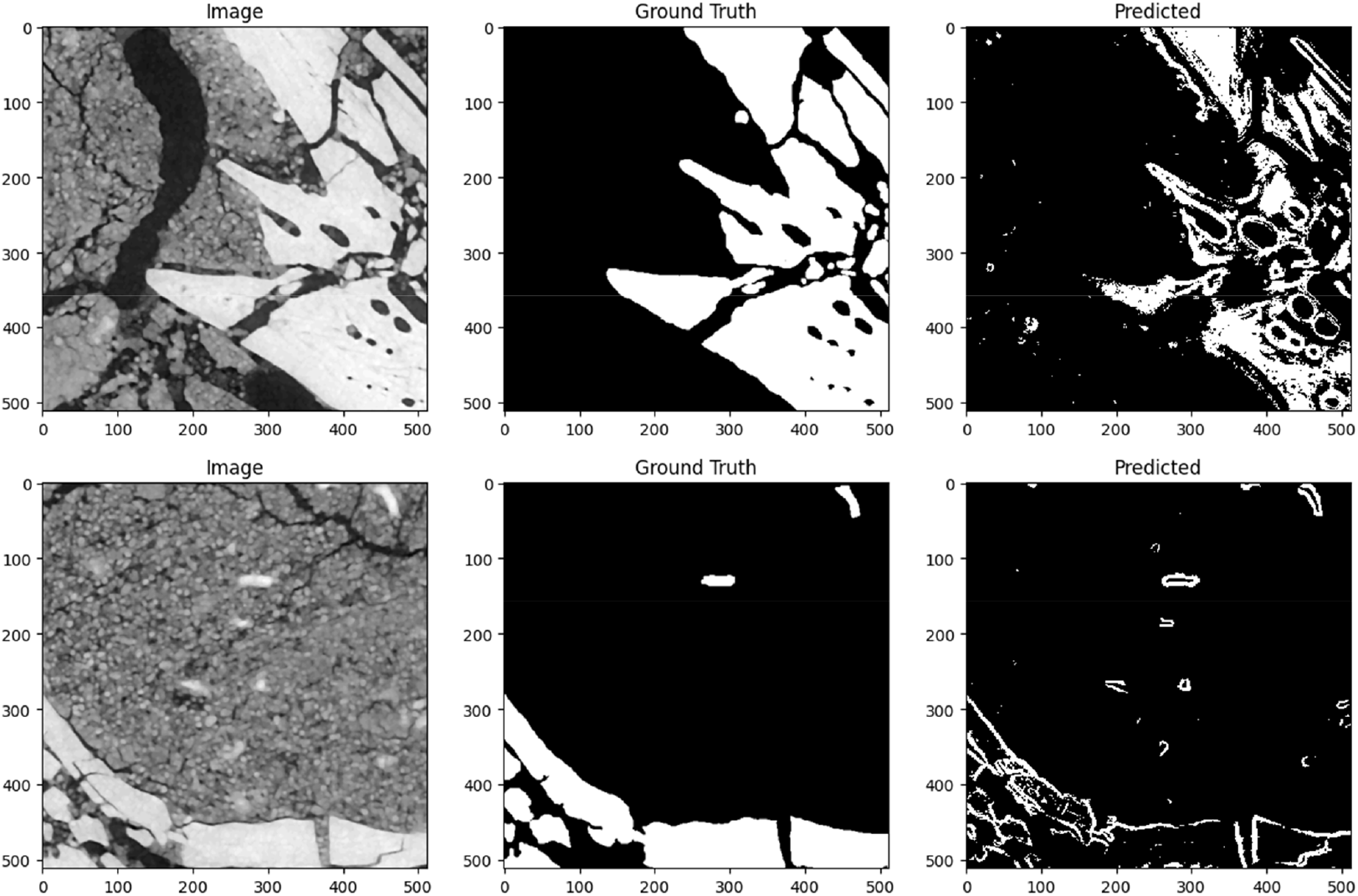
Paddlefish ∼13*µ*m segmentation samples. While the segmentation tends to include the right gross morphology, the model notably failed to segment the internal volume of bones correctly. This interestingly has the effect of producing visually “correct” meshes, since the volume of the segmentation is irrelevant to the surface of the generated OBJ or STL meshes.

Paddlefish ∼45*µ*m Dataset (**Fig. 8**): Performance on this lower-resolution scan was weaker overall, with precision (0.718) and recall (0.191), resulting in an F1-score of 0.185. However, the model achieved high accuracy (0.979), suggesting it effectively avoided false positives, though at the cost of lower sensitivity.

**Fig. 8.**
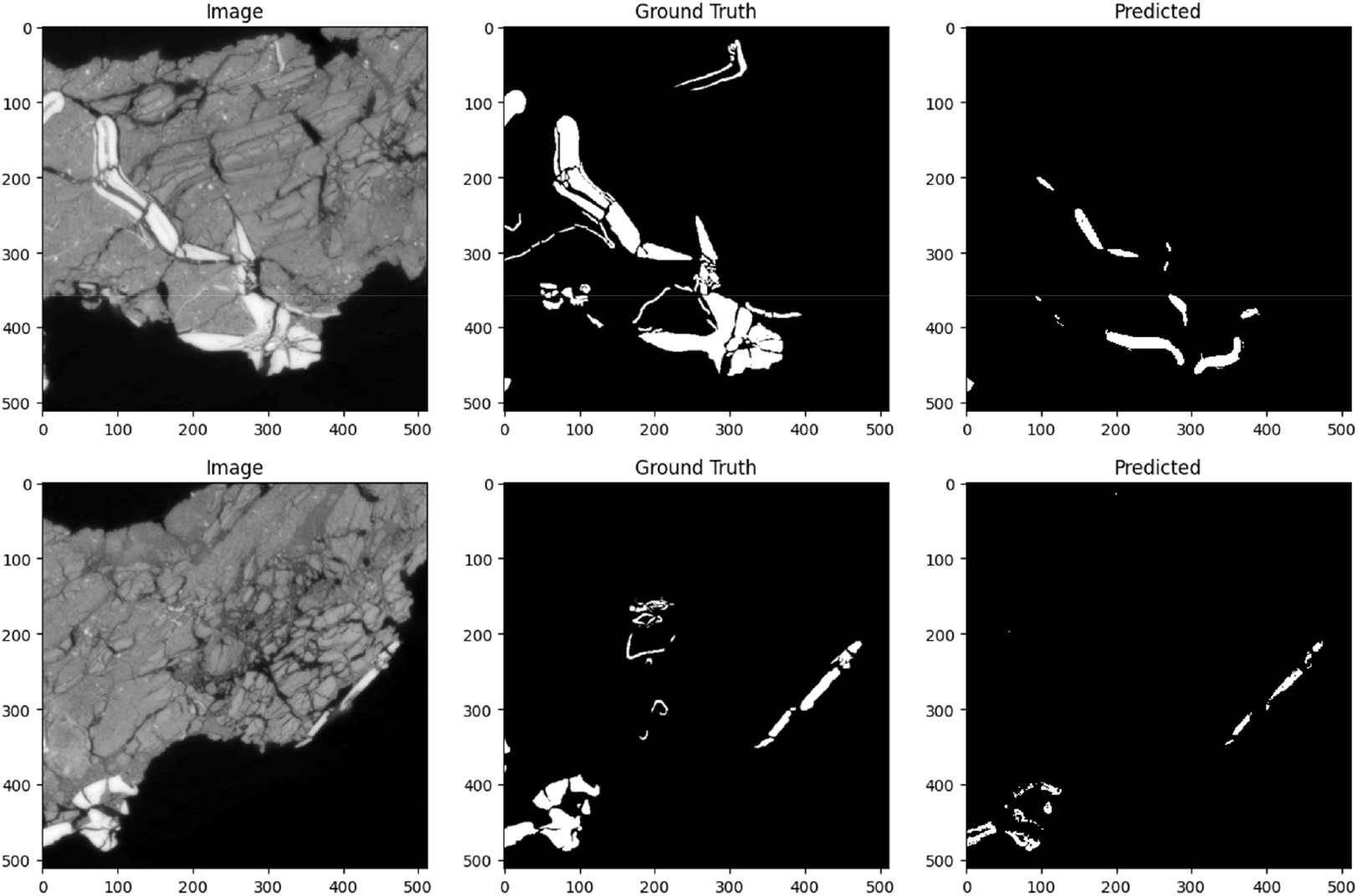
Paddlefish ∼45*µ*m segmentation samples. The model tended to confuse bone for other high-brightness objects in the scene, likely due to underparametrisation and an inability to distinguish bone from other bright sources using the necessary 3D context.

### C. Nothosaur, compared to manual segmentation

The model’s performance declined significantly on this dataset, achieving precision (0.602) and recall (0.496). The low recall suggests that the model struggled to capture true positives. Despite this, accuracy remained reasonably high, likely due to the background dominating the images (**Fig. 9**).

**Fig. 9.**
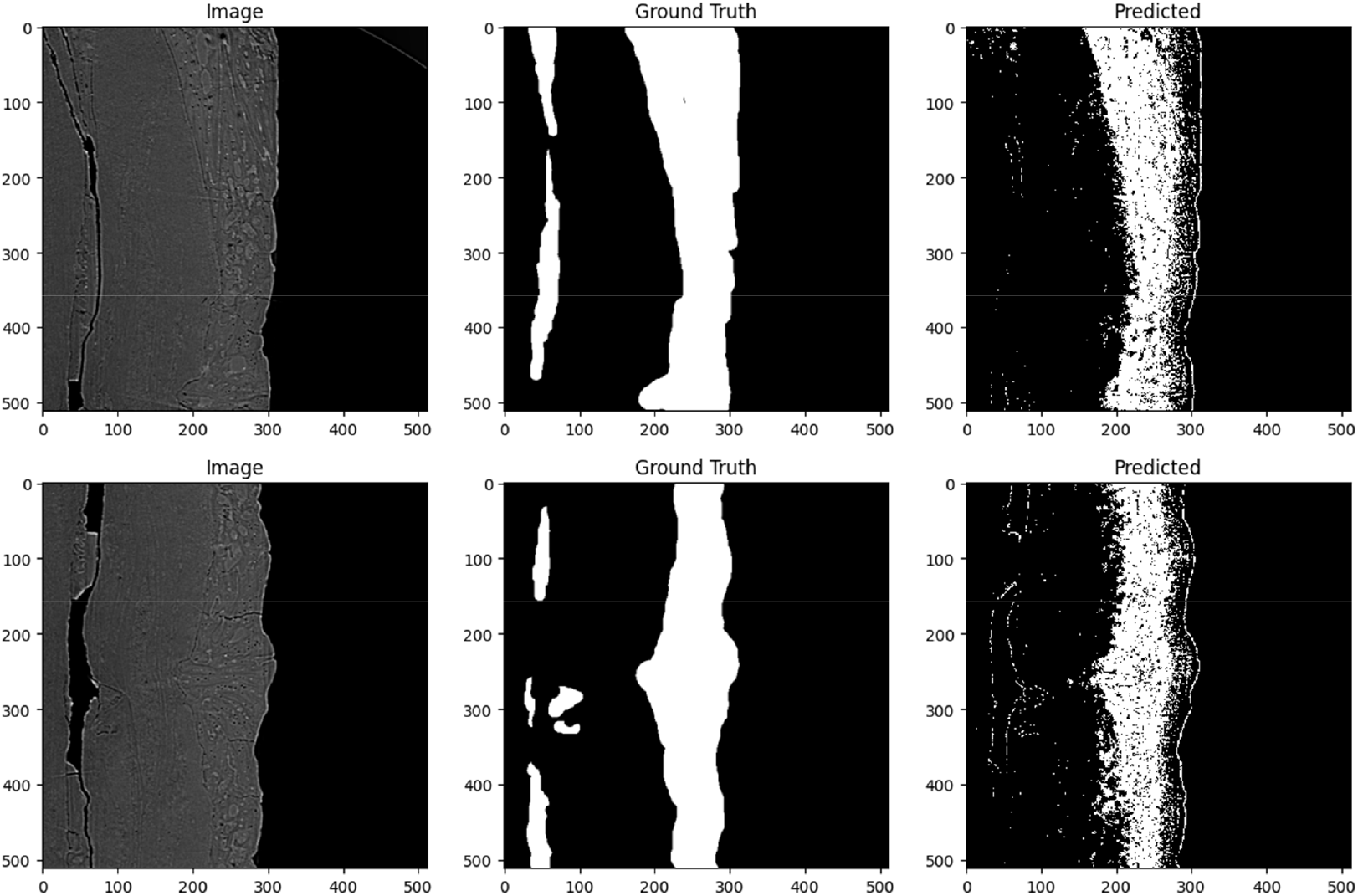
Nothosaur segmentation samples. Two samples of the Nothosaur dataset, illustrating that the texture of the imagery (left panels) was beyond the capabilities of a simple random forest.

### D. Phu Phok Egg data, compared to manual segmentation

This dataset posed the greatest challenge, with extremely low precision which was improved by annotating more slices (n=20 0.232, n=35 0.401) and recall (n=20 0.06, n=35 0.02). Visually, the segmentation quality by the simple random forest was poor overall, reflecting the difficulty encountered by human annotators alike (**Fig. 10**).

**Fig. 10.**
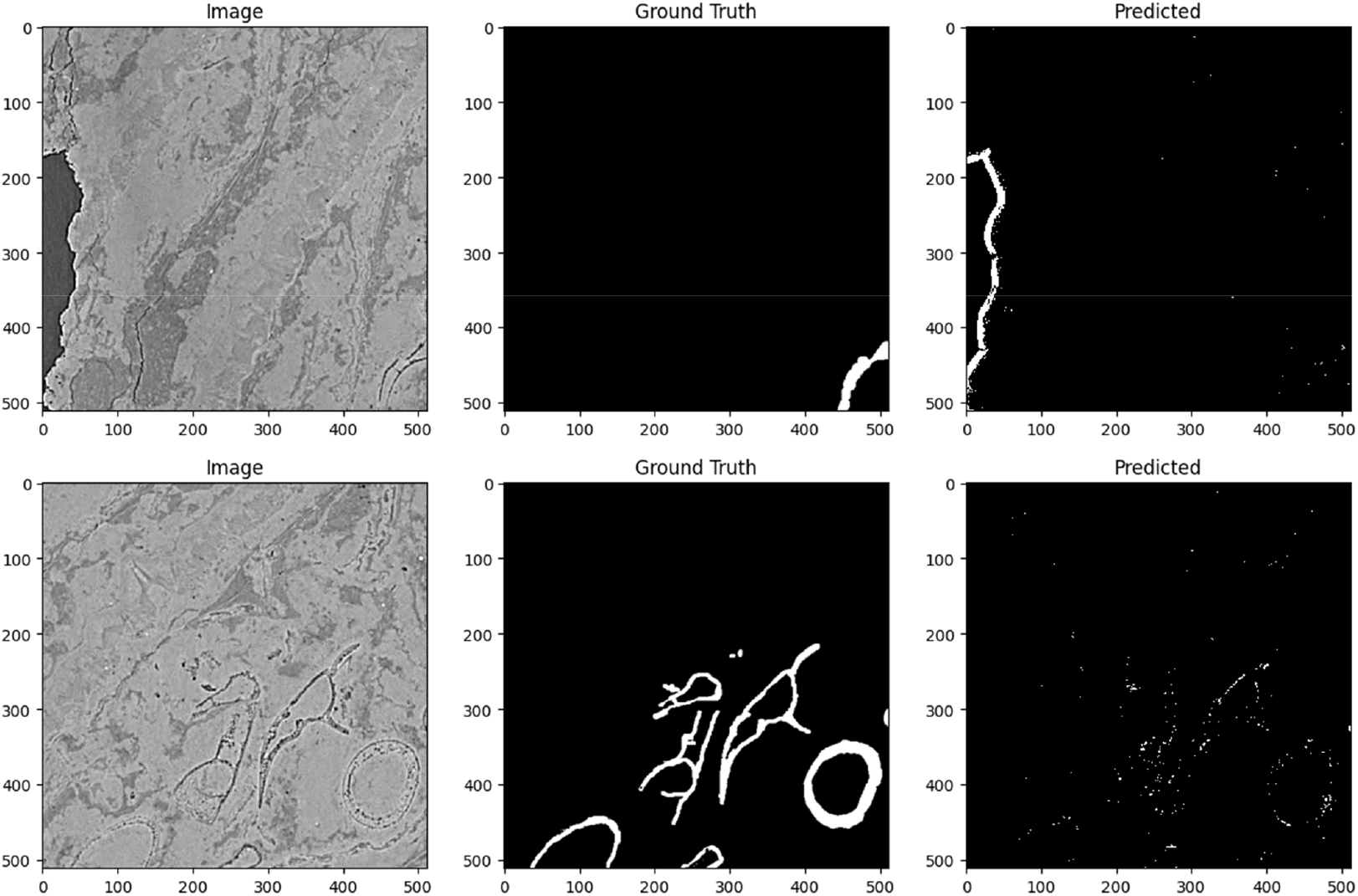
Phu Phok Egg segmentation samples. This challenging dataset illustrates a major failure of the default random forest segmentation model with the multiscale basic pixelwise features in 2D. The high phase contrast in the scan likely caused the model to excessively interpret areas of strong phase contrast as bone, leading it to incorrectly identify other high-contrast edges, such as the boundary of the sample, rather than the low-contrast bones.

## IV. Discussion

The introduction of automated segmentation tools for fossil imaging provides an opportunity to overcome significant bottlenecks in the analysis of large tomographic datasets. As noted in the introduction, manual segmentation is labour-intensive, often taking months or years to complete, which delays scientific advancement. The *ml4paleo* tool offers a valuable contribution by democratising access to segmentation technology, making it accessible to research groups that lack the resources for commercial software solutions.

The results of this study demonstrate that the tool performs reliably on datasets with clear contrast and well-defined features, such as the Burrow dataset, where human-level performance was achieved. The high scores in this case indicate that the model effectively identifies and segments relevant structures. The strong performance on the Paddlefish ∼13µm dataset, also highlights the tool’s capability to handle high-resolution scans with complex anatomical features with a dramatic labour improvement over human annotation alone. However, the drop in performance on the Paddlefish ∼45µm dataset shows that lower-resolution data pose challenges.

The Nothosaur and Phu Phok Egg datasets further highlight the tool’s limitations, particularly when dealing with both low contrast as well as a combination of high phase-contrast with complex low-contrast textures. The segmentation models struggled significantly on both datasets. In the case of the Nothosaur dataset, it is worth noting that obtaining a manual segmentation required over six weeks of full-time effort from a human expert. This underscores the extreme difficulty of the task and puts our tool’s poor performance into perspective, as even manual segmentation was highly labour-intensive due to poor specimen quality. Similarly, segmenting the Phu Phok Egg dataset took approximately eight weeks (pers. comm. Vincent Fernandez), while the Paddlefish scans were segmented in two weeks, of which the majority of time was spent on the ∼45µm dataset. The burrow dataset, however, was segmented quite quickly using simple pixel value thresholding, a process that involves selecting an intensity value and classifying pixels below this threshold as false and those above it as true. All manual segmentations we compared to were performed in VGStudio MAX (volume Graphics, Heidelberg, Germany) and with substantial computing power costs. These results highlight the need for further refinement of the tool and addition of more sophisticated segmentation models, particularly in its ability to handle more challenging datasets with low contrast and intricate textures.

The performance variations across datasets suggest that future versions of the tool could also benefit from specialised models tailored to different fossil types, resolutions, and scan characteristics. Although the current default segmentation algorithm has limitations, the tool’s flexible design allows for the future integration of advanced techniques, such as multi-class segmentation or 3D modelling, to improve its handling of complex fossil data and enhance accuracy across diverse datasets.

More broadly, the *ml4paleo* tool is intended to make fossil segmentation more efficient and accessible. However, segmentation quality can vary across datasets. To mitigate this, the tool supports the replacement or updating of segmentation algorithms, offering users flexibility and enabling ongoing improvements for more consistent results. Additionally, there is potential to enhance the tool’s ability to distinguish closely related structures (e.g., bone and matrix) by incorporating multi-class segmentation and deep learning methods.

We intend for this work to serve as a launch point for future work in progressing the accessibility of palaeontology data and analysis, and that this open-source ecosystem can continue to advance the democratization of science in palaeontology and beyond.

## V. Data & Code Availability

All of our code is open-source and licenced under the Apache 2.0 licence. The code can be accessed at https://github.com/j6k4m8/ml4paleo.

All datasets used in this paper are available through the ESRF Palaeontology database (paleo.esrf.eu). A public version of this tool will be made available upon publication.

## VI. Acknowledgements

We gratefully acknowledge our colleagues for their valuable input in shaping the functional requirements of the tool. We also extend our thanks to Vincent Fernandez for providing his segmentation data and for sharing information regarding the time required to complete these segmentations. M.A.D.D. and P.E.A. were supported by the Swedish Research Council (VR) under grant 2020-03685. F.K.G. and T.B.S. were supported by Kjell och Märta Beijer Foundation. D.F.A.E. was supported by the Dutch Research Council (NWO) under grant 333.22.013. We would also like to thank Dave Marshall for his podcast *Palaeocast*, through which JM discovered the work of MD and colleagues, leading to the initiation of this collaboration.

